# An inducible explant model for dissecting osteoclast-osteoblast coordination in health and disease

**DOI:** 10.1101/2022.10.27.514052

**Authors:** Jarred M. Whitlock, Luis F. de Castro, Michael T. Collins, Leonid V. Chernomordik, Alison M. Boyce

## Abstract

Metabolic bone diseases are a collection of disorders resulting in diminished skeletal integrity and changes in bone mass due to perturbations in the life-long process of bone remodeling. Perturbations in the number, size and nuclear multiplicity of osteoclasts underpin the development of diverse metabolic bone diseases that impact >13% of adults over age 50 world-wide. Each metabolic bone disease (e.g., osteoporosis, Paget’s disease, fibrous dysplasia (FD), osteopetrosis) presents with unique phenotypes, rises from distinct etiologies and progresses with disparate severities, but all are underpinned by a breakdown in osteoclast formation/function. These perturbations of osteoclast formation/function either stem from or cause dysfunctional osteoclast-osteoblast coordination. Unfortunately, a mechanistic understanding of osteoclast-osteoblast coordination and communication is lacking and represents a major barrier to understanding the biology underpinning bone remodeling and the development of effective treatments targeting this process. Here we have developed an inducible *ex vivo* culture model that models osteoclast-osteoblast coordination in the bone remodeling compartment. Doxycycline addition to cultures activates Gα_s_^R201C^ expression and RANKL release from osteoprogenitors, which elicits the differentiation and fusion of neighboring preosteoclasts. In turn, multinucleated osteoclast formation promotes the proliferation of osteoprogenitors, accompanied by the robust release of RANK^+^ extracellular vesicles, all within ∼4 days. This system recapitulates many aspects of the complex osteoclast-osteoblast coordination required for the function of the bone remodeling compartment in both health and diseases underpinned by excessive osteoclast formation. Moreover, based on the ease of isolation, culture, reproducibility and the general adaptability of these cultures to a variety of assays, we expect that this new model will expedite the investigation of osteoclast-osteoblast coordination and osteoclast fusion in bone remodeling and offer a powerful tool for evaluating signaling cascades and novel therapeutic interventions in osteoclast-linked skeletal disease.

**One-Sentence Summary:** Conditional, inducible, *ex vivo* marrow explants offer a novel tool for studying osteoclast formation and osteoclast-osteoblast coordination in a rapid convenient culture model.

## INTRODUCTION

Metabolic bone diseases are a collection of disorders resulting in diminished skeletal integrity and changes in bone mass due to perturbations in the life-long process of bone remodeling (1). Throughout life, the skeleton relies on a basic multicellular unit (BMU) that forms a bone remodeling niche that proceeds along the skeletal matrix and continually remodels bone (2). This BMU is primarily composed of a bipartite remodeling system (i.e., osteoclasts and osteoblasts) and two regulatory cell types that come from the osteoblast lineage and work to orchestrate remodeling through signaling cascades (i.e., bone lining cells and osteocytes). The remodeling system is primarily led by osteoclasts, which erode the inorganic, mineral phase of old/damaged bone and degrade the organic, collagen-rich phase through the release of proteases and osteoblasts, which secrete osteoid and mineralize it to form new bone in its place. In this bipartite remodeling machine, osteoblast lineage cells release a variety of signals that regulate the formation of osteoclasts (3, 4). In turn, recent evidence demonstrates that osteoclasts package and release microRNA rich extracellular vesicles that regulate the differentiation/proliferation of osteoblast precursors (osteoprogenitors) (5-7). Together, the BMU continually erodes old or damaged bone and deposits new to renew the skeletal system (8). Here we offer a novel, rapid model system for resolving how key remodeling members of the BMU – osteoblasts and osteoclasts - coordinate their activities and malfunction with age and disease.

Osteoblast lineage cells activate the formation of multinucleated osteoclasts from mononucleated osteoclast precursors (preosteoclasts), of the myeloid lineage, through the release of osteoclastogenic cytokines (e.g., receptor activator of NF-κB ligand (RANKL)). RANKL binding to the preosteoclast membrane receptor RANK is both necessary and sufficient to upregulate the transcription factors *Nfatc1* and *cFos* and commit precursors to an osteoclast fate (9). Committed osteoclasts then migrate, adhere to one another and fuse plasma membranes (PMs) producing multinucleated, syncytial osteoclasts that typically reach 5-12 nuclei/cell in humans (10, 11), although osteoclasts sizes can differ considerably between individuals and likely increase with age (12, 13). Osteoclast number and size are tightly regulated during bone remodeling, and the number of fusion events forming a multinucleated osteoclast (i.e., its nuclear multiplicity) directly correlates with its ability to resorb bone (i.e., higher nuclear multiplicity = higher bone resorption) (13-16). In addition to eliciting osteoclast formation, osteoblast lineage cells also negatively regulate the number and size of osteoclasts by releasing the RANKL decoy receptor osteoprotegerin (OPG) (17). Thus, tight osteoblast-osteoclast coordination within the bone remodeling compartment manages the number, size, differentiation state and biological activities of this bipartite bone remodeling machine.

Perturbations in the number, size and nuclear multiplicity of osteoclasts underpin the development of diverse metabolic bone diseases that impact >13% of adults over 50 world-wide (>20% of women) (18). Each metabolic bone disease (e.g., osteoporosis, Paget’s disease, fibrous dysplasia (FD), osteopetrosis) presents with unique phenotypes, rises from distinct etiologies and progresses with disparate severities, but all are underpinned by a breakdown in osteoclast formation/function (19-22). These perturbations of osteoclast formation/function either stem from or cause dysfunctional osteoclast-osteoblast coordination. To date, we still lack a detailed understanding of how osteoclasts and osteoblasts coordinate their functions in skeletal biology.

Elucidating a basic blueprint of osteoclast-osteoblast coordination in healthy skeletal remodeling and characterizing how the systems that coordinate this bipartite machine breakdown with age and disease is an essential first step in addressing the growing skeletal health problem in our aging population. Unfortunately, most common models of osteoclast/osteoblast function do not recapitulate the bipartite nature of the bone remodeling machine, but rather recapitulate the isolated function of a single cell type. Immortalized preosteoclast and osteoprogenitor cell lines (i.e., RAW 264.7 and MC3T3-E1 subclones, respectively) are convenient workhorses that suffer from well documented limitations in recapitulating skeletal biology (23-25). Primary osteoclast formation is modeled *in vitro* via enriching primary monocytes that are differentiated stepwise into preosteoclasts and then fusogenic osteoclasts via activation by recombinant macrophage colony stimulating factor (M-CSF) and RANKL (26). Unfortunately, these approaches suffer from the difficulty of obtaining fresh, primary monocytes, the inconsistent propensity of monocytes from different individuals to differentiate and fuse *in vitro*, the significant expense of recombinant M-CSF and RANKL and the inability to passage these cells (13, 27). Obtaining primary osteoprogenitors with consistent, reliable behavior is even more challenging by comparison, requiring the investigator to obtain biopsies from donors or undergo challenging protocols for isolating, enriching and culturing primary cells from murine sources (28, 29). In contrast, animal models do recapitulate the bipartite nature of the BMU and offer *in vivo* context for the underlying biology being evaluated, but introduce significantly longer timelines for experiments; often require specialized equipment for sectioning, demineralizing and imaging cells on bone; introduce greater challenges for collecting high resolution image data; and generally introduce greater complexity and diminished resolution by the nature of osteoclasts and osteoblasts being minor components of the population of cell types that occupy bone.

Here we have developed an inducible explant culture model that recapitulates osteoblast-to-osteoclast signaling and coordination in the BMU. Fibrous dysplasia (FD) is a metabolic bone disease caused by somatic, mosaic gain-of-function mutations in *GNAS* within osteoblast lineage cells, leading to aberrant cell proliferation, suppressed osteoprogenitor differentiation/commitment and the excessive release of osteoclastogenic factors, including RANKL and IL-6 (30-32). We utilized an inducible murine model of FD to develop a tractable *ex vivo* culture model of osteoprogenitor-to-preosteoclast signaling and osteoclastogenesis. Whole bone marrow isolates from the hind limbs of this FD mouse produce a complex cell culture system enriched in osteoprogenitors and preosteoclasts. Doxycycline (Dox.) addition to these complex cultures activates Gα_s_^R201C^ expression and RANKL and IL-6 release from osteoprogenitors, which elicits the differentiation and fusion of neighboring preosteoclasts. In turn, multinucleated osteoclasts release RANK^+^ extracellular vesicles, recently recognized mediators of osteoblast-to-osteoblast signaling, that correlate with changes in the proliferation and differentiation of osteoprogenitors. Moreover, these dynamic biological events all occur within 4 days of Dox. induction. This system recapitulates many aspects of the complex osteoclast-osteoblast coordination required for the faithful function of the BMU that maintains skeletal integrity in health and fails in diseases. Moreover, based on the ease of isolation, culture, reproducibility and the general adaptability of these cultures to a variety of assays, we expect that this new model will expedite the investigation of osteoclast-osteoblast coordination in bone remodeling and offer a simple, first-step tool for evaluating signaling cascades and novel therapeutic interventions in osteoclast-linked skeletal disease.

## RESULTS

### Inducible Gα_s_^R201C^ in osteoprogenitors faithfully elicits osteoclast formation in murine marrow explants

We isolated bone marrow from the tibia and femur of a previously described tet-on Gα_s_^R201C^ murine model, where conditional expression of Gα_s_^R201C^ is specifically induced in osteoprogenitors upon administration of Dox. (22). The tibia and femur from freshly euthanized mice were dissected from both hindlimbs. The marrow space was opened by drilling small holes in the epiphyses with hypodermic needles, whole marrow was flushed out using serum free media and triturated until a homogenous mixture of cells was obtained. These non-adherent isolates were maintained in complete explant media until the majority of cells began to adhere to the culture plastic (∼14 days), whereupon cultures were expanded and passaged for the downstream experiments described below (Fig. 1a). Induction of marrow explant cultures via Dox. activates *Gα*_*s*_^*R201C*^ and *β-gal* transgenes specifically in osteoprogenitors from the skeletal stem cell linage leading to an increase in the steady-state level of Gα_s_ (Fig. 1b). Dox. induction of *Gα*_*s*_^*R201C*^ led to the release of RANKL and IsL-6 from osteoprogenitors and the rapid formation of multinucleated osteoclasts (Fig. 1c-e). β-galactosidase activity observed via x-gal staining provided a linage tracer demonstrating that transgene activation occurs only in osteoprogenitors and not in TRAP^+^ osteoclasts (Fig. 1e). Moreover, the formation of TRAP^+^, multinucleated osteoclasts was dependent on the RANKL produced by Dox. induction of osteoprogenitors, as addition of OPG blocked both the production of TRAP and the observation of multinucleated osteoclasts in induced cultures (Fig. 1e.).

**Figure 1:**
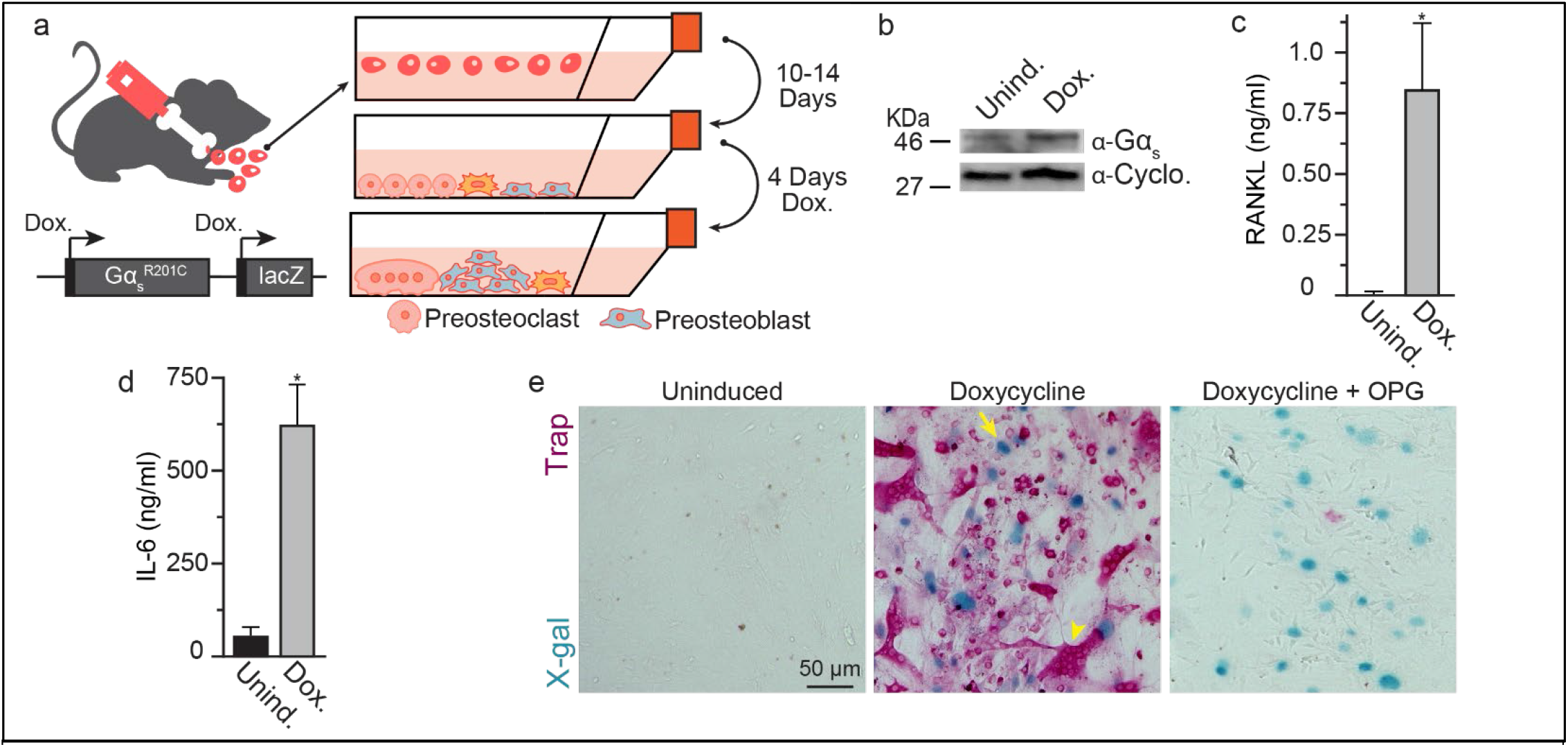
Induction of Gα_s_^R201C^ in osteoprogenitors induces osteoclast formation in marrow cultures. **(a)** A cartoon schematic of our approach for modeling osteoblast-osteoclast signaling/coordination in the formation of osteoclasts. **(b)** A representative Western blot evaluating Gα_s_ expression with (Dox.) or without (Unind.) 4 days induction. **(c)** ELISA evaluation of soluble RANKL in the culture media of marrow explants with (Dox.) or without (Unind.) 4 days induction. (n=7, p=0.02). **(d)** ELISA evaluation of soluble IL-6 in the culture media of marrow explants with (Dox.) or without (Unind.) 4 days induction. (n=4, p=0.01). Representative color brightfield images of marrow explants from Gα_s_^R201C^ following 4 days with (Dox.), without (Unind.) or with induction in the presence of OPG (Dox.+OPG). Arrow denotes x-gal staining of the nucleus of an induced osteoprogenitor, Arrowhead denotes a multinucleated, TRAP^+^ osteoclast. Error bars=SEM. Significance was assessed via paired t-test.

### Osteoprogenitor signaling elicits robust osteoclastogenesis in a Rankl-dependent manner

Tet-on Gα_s_^R201C^ marrow explant cultures offer a rapid, tractable model for analyzing osteoclast formation elicited by osteoprogenitors in an *ex vivo* environment, modeling the multicellular coordination of the BMU. Dox-induced osteoprogenitor-to-preosteoclast signaling within these cultures activated osteoclastogenesis in preosteoclasts that fuse and form syncytia with typical ruffled borders in ∼4 days (Fig. 2a). We observe negligible multinucleated osteoclast formation prior to 2 days of induction and a near linear increase in osteoclast fusion until 4 days of induction, when fusion plateaued (Fig. 2b). RANKL is essential for activating osteoclast formation from preosteoclasts, as Dox. induction activated the upregulation of the osteoclastogenic transcription factors *Nfatc1* and *cFos*, both of which were blocked by the addition of OPG, the RANKL decoy receptor (Fig. 2c). Upregulation of *Nfatc1* and *cFos* coincides to the formation of TRAP^+^, multinucleated osteoclasts and expression of TRAP and extent of fusion between preosteoclasts depend on RANKL production, as multinucleated cells were not observed in cultures without Dox. induction and OPG suppressed their formation in a concentration dependent manner (Fig. 2d). Osteoclasts not only fused in response to Dox. induction, but the formation of multinucleated osteoclasts was associated with their production of the resorptive enzyme cathepsin K (Fig. 2e). In addition to general cytoskeletal and chromogenic staining, we found that immunofluorescence imaging of the pan-macrophage lineage marker CD68 and the osteoclast enriched sorting nexin Snx10 easily facilitated the detection of preosteoclasts and subsequent multinucleated osteoclasts among the cell types present in our complex cultures (Fig. 2f). Finally, Dox. induction of marrow explant cultures not only elicited the formation of osteoclast syncytia, but also activated their resorptive activity in a RANKL dependent manner (Fig. 2g).

**Figure 2:**
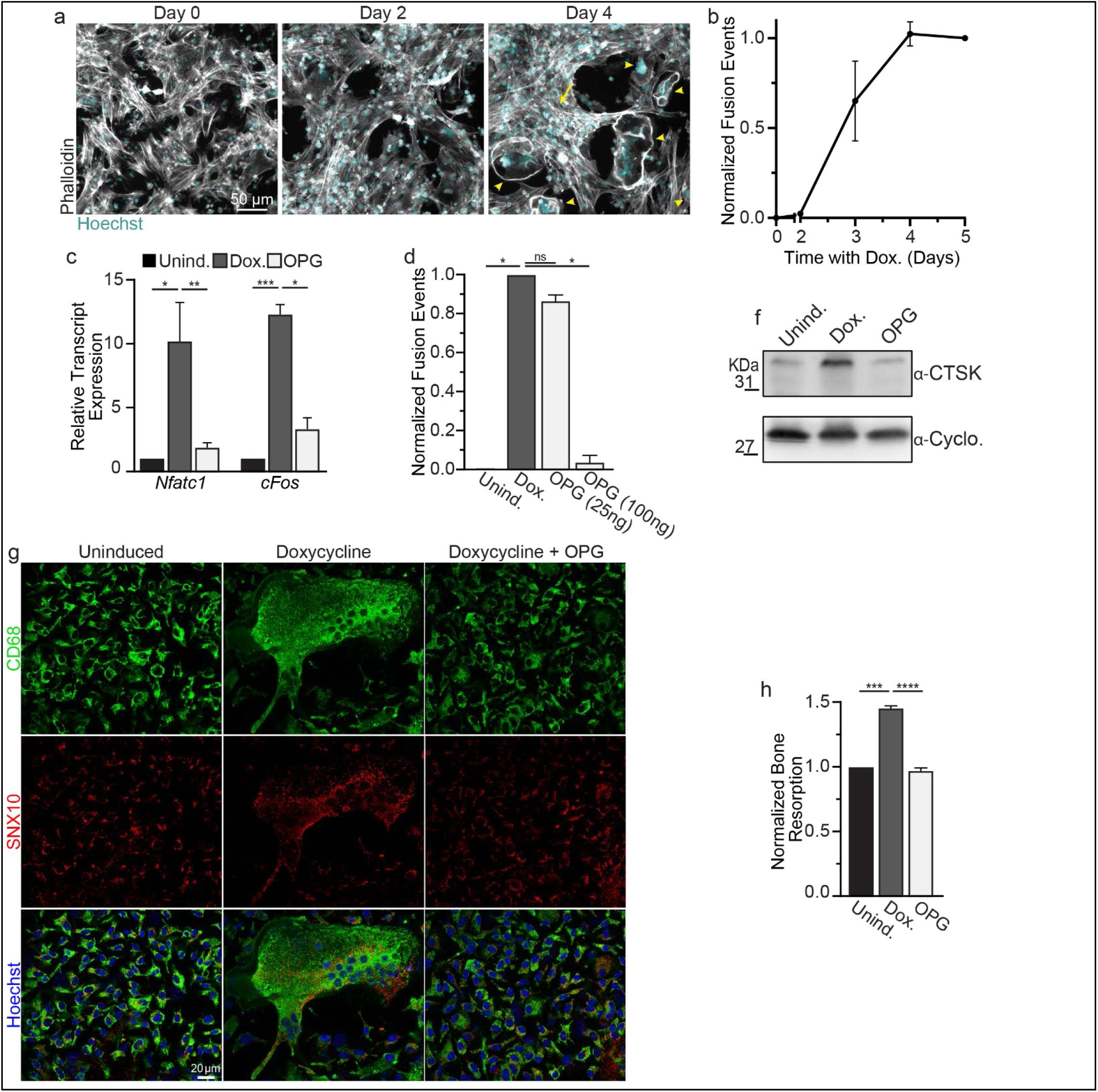
Dox. induction elicits osteoclast formation. **(a)** Representative images of *ex vivo* marrow cultures following Dox. induction (Grey = Phalloidin-Alexa488, Cyan = Hoechst). **(b)** Quantification of the number of fusion events over time following addition of Dox. Each time course was normalized to the number of fusion events observed on day 5. (n=3) **(c)** qPCR evaluation of the osteoclastogenesis transcription factors *Nfatc1* and *cFos* under uninduced (black), Dox. induced (grey) or Dox. Induced + 100ng/ml OPG (white) conditions. Fold expression is relative to *Actb* RNA. (n=3) **(d)** Quantification of the number of fusion events in *ex vivo* marrow cultures treated as in **c. (e)** Representative Western blot evaluation of the steady-state level of the osteoclast resorption enzyme CTSK in *ex vivo* marrow cultures treated as in **c. (f)** Representative immunofluorescence images of the pan macrophage lineage marker CD68 (green), the osteoclast sorting nexin SNX10 and Hoechst in *ex vivo* marrow cultures treated as in **c. (g)** Quantification of bone resorption by *ex vivo* marrow cultures treated as in **c**. In **a** and d, Arrowheads = multinucleated osteoclasts, Arrows = proliferative preosteoblasts. Error bars=SEM. Statistical significance was assessed via paired t-test.

### Osteoclast formation promotes osteoprogenitor proliferation

The formation of multinucleated osteoclasts was also accompanied by the development of fibrous, multicellular lesions, analogous to those observed in patients with gain-of-function mutations in Gα_s_ (Fig. 2a, 3a, arrows). Staining of these lesions demonstrated that the bulk of cells exhibited enriched, nuclear staining for the osteoblast precursor (preosteoblast) marker Runx2 (Fig. 3b). While preosteoblasts were readily observed in uninduced cultures, neither enrichment nor the clumping phenomenon was observed in the absence of Dox. induction (Fig. 3a). Further underscoring the proliferative nature of these multicellular lesions, the preosteoblasts demonstrated a dramatic nuclear enrichment of the proliferative marker Ki67 (Fig. 3b-c). Moreover, in addition to the increased proliferation observed in preosteoblasts, we observed a marked decrease in the osteoblast differentiation markers *Sp7, Dxl5* and *Bglap* (33) in our cultures following Dox. induction, suggesting reduced osteogenic commitment accompanied this increase in preosteoblast proliferation (Fig. 3d). As with the proliferative phenotype, the marked decrease in osteoblast differentiation observed was partially rescued by blocking the formation of osteoclasts via OPG ablation of osteoclasts, suggesting that osteoclasts contribute to the observed changes in preosteoblast proliferation/differentiation (Fig. 3c-d).

**Figure 3:**
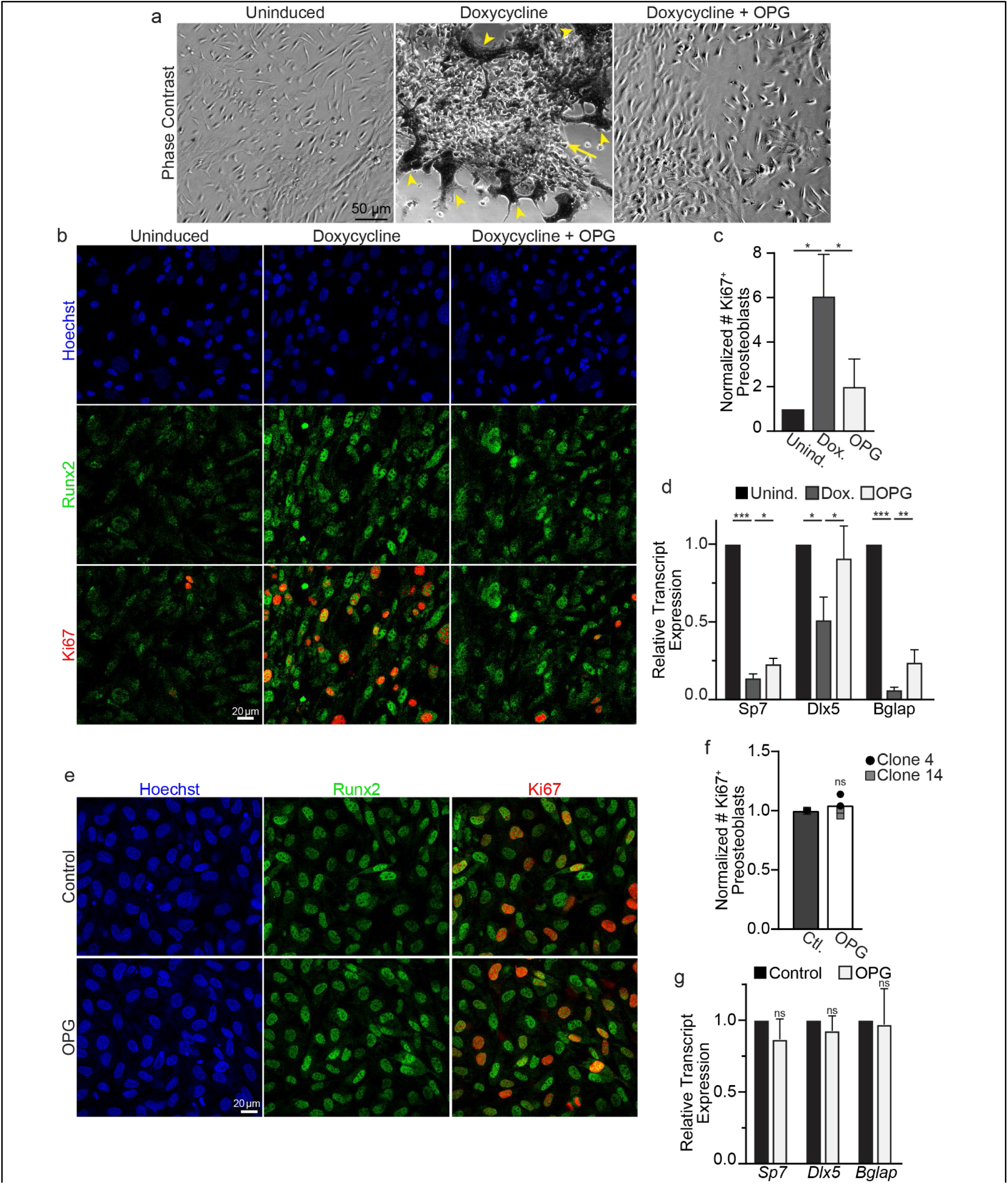
Osteoclast formation in *ex vivo* cultures elicits proliferation in osteoprogenitors. **(a)** Representative phase contrast images of TRAP stained marrow explants from a Gα_s_^R201C^ following 4 days with (Dox.), without (Unind.) or with induction in the presence of OPG (Dox.+OPG). Arrow denotes lesion of cells, Arrowheads denote TRAP^+^, multinucleated osteoclasts, which appear dark in phase contrast. **(b)** Representative immunofluorescence images of preosteoblast marker Runx2, proliferation marker Ki67 and Hoechst in cultures treated as in **a. (c)** Quantification of the fraction of Runx2^+^ preosteoblasts that exhibit staining for the proliferation marker Ki67. (n=3) **(d)** qPCR evaluation of the osteogenic markers *Sp7, Dlx5* and *Bglap* under uninduced (black), Dox. induced (grey) or Dox. induced + 100ng/ml OPG (white) conditions. Fold expression is relative to *18s* rRNA. (n=5) **(e)** Representative immunofluorescence images of preosteoblast marker Runx2, proliferation marker Ki67 and Hoechst in MC3T3-E1 clone 14 preosteoblasts treated with (OPG) or without (control) 100ng/ml OPG. **(f)** Quantification of the fraction of Runx2^+^ preosteoblasts that exhibit staining for the proliferation marker Ki67 treated as in **e**. (n=3 for clone 4(circle), n=3 for clone 14(square). Bar denotes summary average of both clones 4 and 14) **(g)** qPCR evaluation of the osteogenic markers *Sp7, Dlx5* and *Bglap* in clone 14 under control (black), 100ng/ml OPG (white) conditions. Fold expression is relative to *18s* rRNA. (n=3) Error bars=SEM. Significance was assessed via paired t-test.

In addition to releasing soluble RANKL, preosteoblasts express a transmembrane RANKL variant. To test whether changes in preosteoblast proliferation/differentiation elicited by OPG may be caused by some direct effect OPG has on preosteoblasts, we treated two preosteoblast cell lines (MC3T3-E1 clone 4 and clone 14) with OPG and looked for changes in Ki67 staining. In both clones, we observed no change in the expression of Runx2 or Ki67 staining, suggesting that the effect of OPG addition on the proliferation of preosteoblasts in our explant cultures is not direct (Fig. 3e-f). Likewise, the addition of OPG to preosteoblasts alone had no impact on the steady-state levels of the osteoblast differentiation marks observed in our *ex vivo* cultures (Fig. 3g vs 3d). Together, these data suggest that proliferative/differentiation changes in the preosteoblasts in our explant model are mediated by osteoclasts and are not a direct effect of OPG on preosteoblasts.

### Osteoprogenitor proliferation correlates with the release of Rank^+^ extracellular vesicles (EVs)

Recent reports have highlighted the role of EVs as mediators of osteoclast-to-osteoprogenitor signaling and regulation in coordinated bone remodeling (5, 33-35). Previous characterizations of osteoclast EVs have identified both RANK and TGS101 as markers of osteoclast derived EVs that impact osteoclast and osteoblast differentiation (5-7, 36). To assess whether our system models the osteoclast production of EVs that coordinate osteoclast-osteoblast activity, we collected conditioned media from our cultures and evaluated whether they contained EVs released by osteoclasts. We induced our cultures and waited until we observed multinucleated osteoclasts (3-4 days). Next, we placed our cultures in complete explant media made with EV depleted FBS, allowed cultures to condition the media for 24 hours, collected the media and enriched an EV fraction using a simple centrifugation approach (Fig. 4a). Biochemical assessment of these EV-enriched fractions demonstrated that Dox. induction led to an enrichment of the osteoclast transmembrane receptor RANK and the exosome marker TSG101 in the EV fraction (Fig. 4a). The enrichment of RANK and TSG101 in the EV fraction produced by explant cultures was not observed without Dox. induction and was suppressed by OPG inhibition of osteoclast formation (Fig. 4a-c). Identical results were also obtained using commercially available EV isolation kits used according to manufactures instructions (QIAGEN ExoEasy). These findings indicated that osteoprogenitor-dependent osteoclast formation in our model is accompanied by the release of osteoclast-generated, RANK^+^ EVs.

**Figure 4:**
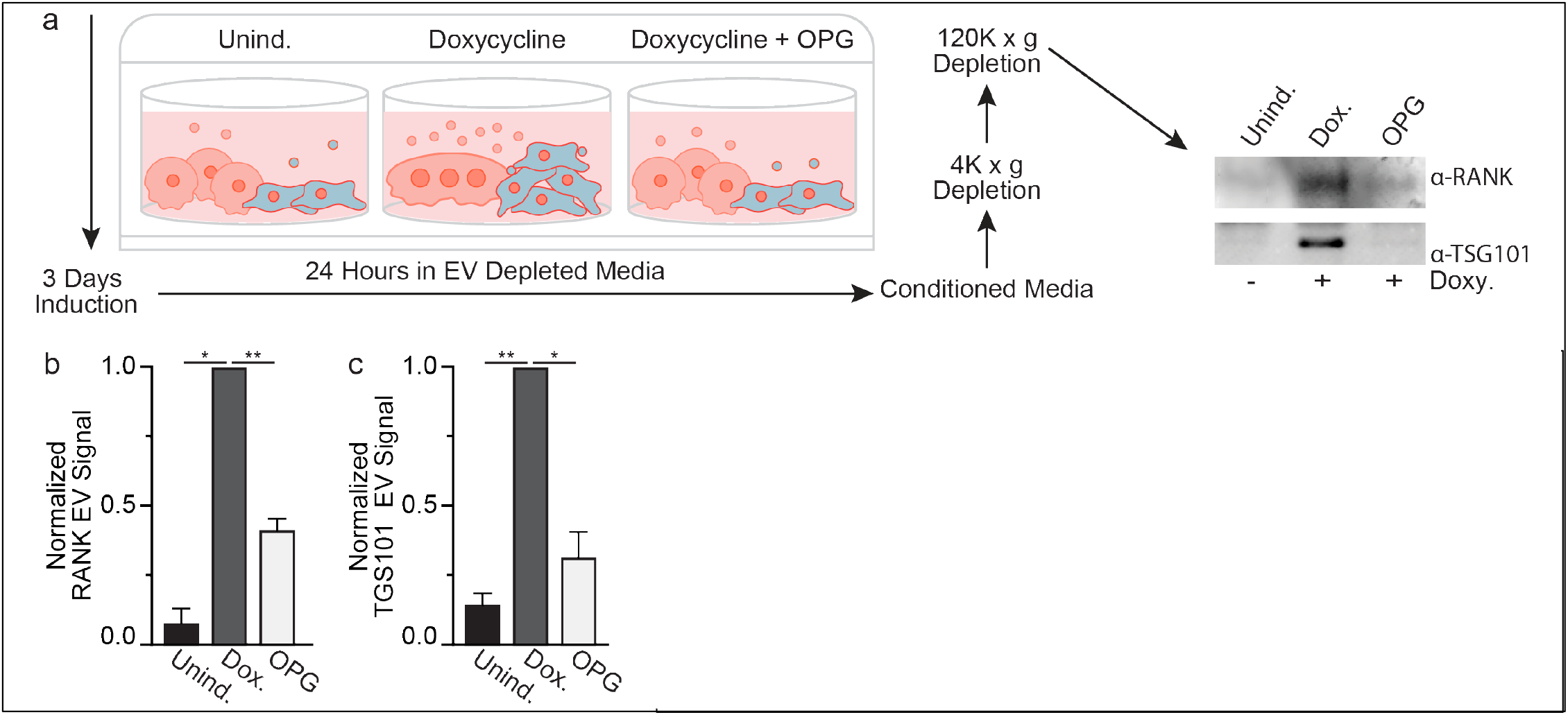
Osteoclast formation in *ex vivo* marrow cultures is accompanied by EV release. **(a)** An illustrated schematic depicting an approach for enriching an EV enriched fraction from uninduced, Doxy. induced and Doxy. induced + 100 ng/ml OPG *ex vivo* marrow cultures (left). A representative Western blot evaluating EV enriched fractions for the osteoclast membrane receptor RANK and the exosome marker TSG101 (right). **(b)** Quantification of RANK signal in Western blots of EV enriched fractions, as in **a**. (n=3) **(c)** Quantification of TSG101 signal in Western blots of EV enriched fractions, as in **a**. (n=5) Error bars=SEM. Significance was assessed via paired t-test.

In summary, we have established a simple, inducible, tractable tool for the study of osteoprogenitor induced osteoclast formation that can be implemented to study how osteoclast and osteoblast lineage cells coordinate their functions in health and disease.

## DISCUSSION

Here we describe a primary culture model for recapitulating the dynamic interactions and regulatory relationships between the primary cellular components of the bipartite bone remodeling machine. Although the cellular make-up of the explant cultures described and the biological activities represented are complex, employing this model is relatively simple. The training required to master the simple dissections required is minimal, the reproducibility between cultures and passages is outstanding and the cost in maintaining and implementing this model of osteoclast/osteoprogenitor biology is significantly less than even the simplest cellular models of osteoclast biology, as the only media additive required for activation is Dox.

Our primary aim was to develop a model of osteoclast formation that was activated by biologically relevant stimuli and not the artificial addition of a supraphysiological dose of the osteoclastogenic factor (RANKL). Here we have developed a culture model where osteoprogenitors are specifically induced to overexpress a gain-of-function mutation that elevates cAMP and promotes the release of endogenous osteoclastogenic factors (e.g., RANKL, IL-6, likely others). These factors activate osteoclastogenesis, promote osteoclast fusion, elicit osteoclast production of resorptive enzymes and lead to osteoclast-dependent demineralization in relatively short timeline (4 days following Dox. addition). Moreover, these cultures easily adapt to larger culture ware for biochemical sample collection (T-75 flasks, multi-well plates for screening assays, polymer bottom μ-slides for high-resolution imaging and mineralized biomimetics of bone) with no apparent impact on their ability to faithfully produced osteoprogenitor-dependent multinucleated osteoclasts.

In addition to osteoclasts, we find that osteoblast lineage cells are well represented in the cultures. Skeletal stem cell progeny are the only cells containing the conditional tet-on Gα_s_^R201C^ cassette responsible for RANKL production following Dox. addition, as documented in the murine model from which our cultures are derived (22). Moreover, our addition of a tet-on β-gal cassette to this these cells allows for simple linage tracing and the detection of induced cells. Gα_s_^R201C^ is a common gain-of-function mutation in osteoprogenitors that leads to the excessive production of RANKL and the ectopic osteoclast formation that underpins the pathophysiology of FD. By taking advantage of this FD model, we have developed a tool that facilitates the ability to follow osteoprogenitor-to-osteoclast signaling with high spatial and temporal resolution. It is our hope that future work will further develop our model as a tool for evaluating the propensity of the osteoblast lineage cells in our cultures to secrete matrix and mineralize it. Although we have focused on cells of the osteoprogenitor and preosteoclast lineage, we cannot dismiss the possible presence or contribution of other cell types to the ability to passage, grow and faithfully induce cultures to elicit the preosteoblast-dependent formation of multinucleated osteoclasts. However, after the initial passage of the cell types that adhere to culture plastic, we observe that the vast majority of cells within our cultures exhibit morphology consistent with CD68^+^ macrophages or RUNX2^+^ osteoblast precursors we typically stain in our experiments and that nearly all cells stain positively for either β-gal or TRAP activity following Dox. induction.

In addition to observing changes in osteoprogenitors and osteoclasts, our model offers an excellent opportunity to study the complex cell-cell signaling that coordinates the functions of the BMU during bone remodeling. Well established intercellular signaling cascades tightly regulate the formation and activity of the multicellular unit (e.g., RANKL, IL6, TGF-β, BMPs), but growing evidence suggests that a complex array of cytokines and secondary messengers contribute to the formation of osteoblasts, osteoclasts and their coordination. Moreover, new evidence demonstrates the role of EVs in coordinating osteoclast-osteoblast function and acting as a feedback mechanism for managing biological activity in bone remodeling (5, 35, 36). Recent work has also demonstrated dramatic changes in EV signaling in bone disease and injury, highlighting the clinical need for reliable systems with which to identify the signals relayed in EVs released by osteoclasts/osteoblasts and how therapeutic interventions modulate the release of these critical secondary messengers and their contents (34, 37, 38). Here we demonstrate that our *ex vivo* cultures are rich sources of these EV secondary messengers exchanged by constituents of the BMU. We expect that these cultures will be an excellent tool for studying the contents, biological activity, signaling properties and downstream effects of these and other secondary messengers in the processes of coordinated bone remodeling.

Unfortunately, many current models of osteoclast/osteoblast function only replicate one constituent of the BMU that remodels bone or a complete remodeling compartment *in vivo*. These approaches either suffer from a lack of biological context or come with a level of complexity that makes the high-resolution evaluation of a relatively minor component of bone challenging. Here we describe a conditional, inducible, *ex vivo* approach that facilitates the rapid assessment of osteoprogenitor-preosteoclast coordination, the formation of osteoclasts and osteoclast-to-osteoprogenitor signaling in a fast, highly adaptive, cost-effective culture model (Fig. 5). While our system is built from genetic tools used to model FD, we would like to emphasize that its applications far surpass the study of this single disease. Instead, here we have a multicellular unit where osteoclast formation, activity and signaling is directly inducible under the guidance of an osteoprogenitor conductor. We believe our culture system will be useful for addressing mechanisms underpinning the basic formation and coordination of osteoclasts and osteoblasts, as an acute model of the excessive osteoclast formation that accompanies many resorptive bone diseases and can help us better address how osteoblast-osteoclast coordination is accomplished in health and disease. We have utilized this novel model in two recent manuscripts to identify a novel regulator of osteoclast fusion and assess the molecular mechanism of Denosumab treatment of fibrous dysplasia (39, 40). It is now our hope to disseminate this tool and partner with our colleagues to implement its use in answering new, vital questions concerning how the BMU and bone remodeling are managed in health and disease.

**Figure 5:**
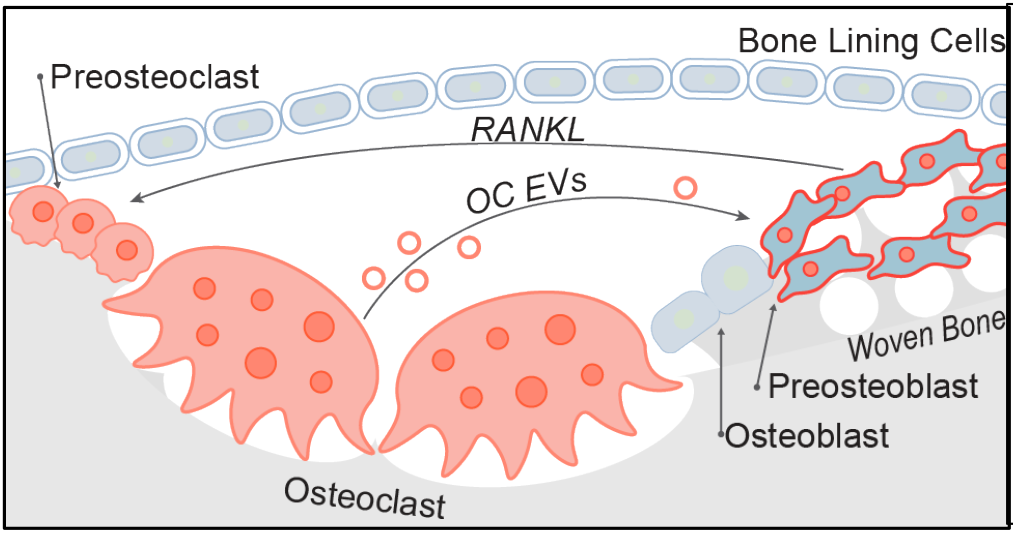
*Ex vivo* FD marrow explants as a model of osteoclast formation and osteoclast-osteoblast coordination. **(a)** An illustration of the biological process modeled in our *ex vivo* marrow cultures. Doxycycline induction elicits preosteoblast expression of Gα_s_^R201C^ and release of RANKL. RANKL binding initiates osteoclastogenesis and the formation of multinucleated, bone resorbing osteoclasts. Osteoclasts release EVs that correlate with preosteoblast proliferation. Peach cells are of the monocyte>osteoclast lineage. Blue cells are of the skeletal stem cell>osteoblast lineage.

## MATERIALS AND METHODS

### Reagents

TRAP staining reagents were purchased from Cosmo Bio Co. (#PMC-AK04F-COS). Bone Resorption Assay Kits were purchased from Cosmo Bio Co. (Catalogue # CSR-BRA-24KIT) and used according to the manufacturer’s instructions. Hoechst 33342 and phalloidin-Alexa555 were purchased from Invitrogen (#H3570 and A30106, respectively). Doxycycline hydrochloride was purchased from Sigma.

### Animals

All animal studies were carried out according to NIH-Intramural Animal Care and Use Committee (ACUC) of the National Institute of Dental and Craniofacial Research approved protocols (ASP #19-897), in compliance with the Guide for the Care and Use of Laboratory Animals. A mouse model of fibrous dysplasia with inducible expression of hyperactive GαsR201C in cells of the osteogenic lineage (41) was used to obtain bone marrow explants (described below). For this study we used 12-18 week old females.

### Murine Bone Marrow Explant Culture

The tibia and femur were dissected from an inducible murine model of fibrous dysplasia described previously (41) or wild-type littermates. Holes were drilled into the epyphises of each bone using a 22-gauge hypodermic needle, and the bone marrow was flushed into a culture dish using alpha MEM. These bone marrow isolates were further dissociated through a fresh 22-gauge hypodermic needle to obtain a single cell suspension, and cultured in alpha MEM plus 20% FBS, 1x pen/strep (complete explant culture media) and 1x Normocin (InvivoGen, # Ant-nr-1) for 12-14 days in T-75 culture flasks. Cells that adhered to the flask were washed 3 times with PBS and passaged using 0.05% Trypsin and a cell scraper and cultured for up to 4 passages in complete explant culture media, less Normocin. For GαsR201C expression induction in the bone marrow stromal cell subset of the explants, cells were plated at ∼40% confluency in 6-well plates, 24-well plates, and angiogenesis µ-slides (Ibidy) for EV collection and staining, respectively. For induction, cells were treated with 1µM doxycycline (Sigma, # D9891-5G). During induction, media were refreshed daily. OPG was added at 100ng/ml.

### Antibodies

We used α-SNX10 (SIGMA, SAB2107086), α-TSG101 (NOVUS, 4A10), α-CD68 (Abcam, 53444), α-RUNX2 (Abcam, 76956), α-Ki-67 (CST, D3B5), RANK (Abcam, 13918), α-Ms-Alexa488 (CST, 4408), α-Ms-Alexa555 (CST, 4409) α-Rb-Alexa488 (CST, 4412), α-Rb-Alexa555 (CST, 4413), and α-Rt-Alexa488 (INV, 1106).

### Fluorescence Microscopy Imaging

In the immunofluorescence experiments we washed the cells with PBS and fixed with warm freshly prepared 4% formaldehyde in PBS (Sigma, F1268) at 37^0^C. The cells were washed three times with PBS. To permeabilize the cells, we incubated them for 10 min in PBS with 0.1% Triton X100 in PBS and 5% FBS (IF Buffer) to suppress non-specific binding. Then the cells were incubated with primary antibodies overnight in IF Buffer. After 5 washes in IF Buffer, we placed the cells for 2h at room temperature in IF Buffer with secondary antibodies. Cells were then again washed 3 times with IF Buffer followed by 2 times with PBS. Images were captured on a Zeiss LSM 800 airyscan, confocal microscope using a C-Apochromat 63x/1.2 water immersion objective.

### Bone Resorption

Bone resorption was evaluated using bone resorption assay kits from Cosmo Bio USA according to the manufacturer’s instructions. In short, fluoresceinamine-labeled chondroitin sulfate was used to label 24-well, calcium phosphate-coated plates. Human, monocyte-derived osteoclasts were differentiated as described above, using alpha MEM without phenol red. Media were collected at 4-5 days post RANKL addition, and fluorescence intensity within the media was evaluated as recommended by the manufacture.

### Fusion Assays

Osteoclast fusion was evaluated by fluorescence microscopy (42). Briefly, cells were fixed with 4% paraformaldehyde at timepoints of interest, permeabilized with 0.1% Triton X-100 and blocked with 5% FBS. Cells were then stained with phallodin-Alexa488 and Hoechst to label cells’ actin cytoskeleton and nuclei, respectively. 8 randomly selected fields of view were imaged using Alexa488, Hoechst and phase contrast compatible filter sets (BioTek) on a Lionheart FX microscope using a 10x/0.3 NA Plan Fluorite WD objective lense (BioTek) using Gen3.10 software (BioTek). Osteoclast fusion efficiency was evaluated as the number of fusion events between osteoclasts with obvious ruffled boarders and ≥3 nuclei in these images, as described previously (43). Since regardless of the sequence of fusion events, the number of cell-to-cell fusion events required to generate syncytium with N nuclei is always equal to N-1, we calculated the fusion number index as Σ (Ni − 1) = Ntotal − Nsyn, where Ni = the number of nuclei in individual syncytia and Nsyn = the total number of syncytia. In contrast to traditional fusion index measurements, this approach gives equal consideration to fusion between two mononucleated cells, one mononucleated cell and one multinucleated cell and two multinucleated cells. In traditional fusion index calculations, fusion between two multinucleated cells does not change the percentage of nuclei in syncytia. If instead one counts the number of syncytia, a fusion event between two multinucleated is not just missed but decreases the number of syncytia. In contrast, the fusion number index is inclusive of all fusion events.

### Transcript Analysis

For real-time PCR, total RNA was collected from cell lysates using PureLink RNA kit following the manufacturer’s instructions (Invitrogen # 12183018A). cDNA was generated from total RNA via reverse transcription reaction using an iScript RNA-to-cDNA kit according to the manufacturer’s instructions (BioRad). cDNA was then amplified using the iQ SYBR Green Supermix (Biorad). All primers were predesigned KiCqStart SYBR Green primers with the highest rank score specific for the gene of interest or GAPDH control and were used according to the manufacturer’s instructions (Sigma). All Real-time PCR reactions were performed and analyzed on a CFX96 real-time system (Biorad), using 18s ribosomal RNA as an internal control. Fold-change of gene expression was determined using the ΔΔCt method (44). 3-4 independent experiments were performed, and each was analyzed in duplicate.

### Enrichment of Extracellular Vesicle Fractions

*Ex vivo* cultures were or were not induced under the conditions described for 4 days and then switched to complete explant culture media made with FBS depleted of EVs (depleted via ultracentrifugation at 150,000xg for >2 hours). Cultures were allowed to condition the media for 24 hours. Conditioned media were collected, and cells/large cell debris were depleted via centrifugation (15 mins at 4,000xg). Next, an EV fraction was enriched via centrifugation (150,000xg for 1.5 hours). EV enriched fractions were evaluated via Western Blot using α-TSG101 the α-RANK antibodies described below. Similar results were also obtained using the exoEasy Maxi Kit (Qiagen) following the manufacture’s instructions. TSG101 and RANK signals were quantified using densitometry via ImageJ.

## Author Contributions

J.M.W., L.F.C.D., and L.V.C. designed the model described and the experiments to implement its use. J.M.W. performed the experiments described, analyzed the data and produced all figures. M.T.C and A.M.B. contributed intellectually to experimental approaches for assessing osteoclast-to-osteoprogenitor crosstalk. L.F.C.D. maintained the mouse model described and carried out genetic crosses to introduce the β-gal lineage tracer and ensure the fidelity of the model. J.M.W. and L.V.C. wrote the manuscript with assistance from L.F.C.D., M.T.C., and A.M.B.

## Acknowledgements

We thank Rebeca Galisteo and Tiffani Slaughter of M.T.C’s laboratory for providing the sacrificed mice from which explant cultures were isolated. The research in L.V.C. laboratory was supported by the Intramural Research Program of the Eunice Kennedy Shriver National Institute of Child Health and Human Development, National Institutes of Health. The research in A.M.B.’ and M.T.C.’ laboratories was supported by the Intramural Research Program of the National Institute of Dental and Craniofacial Research, National Institutes of Health. Work in MTC lab and LVC labs was also supported by the of Research on Women’s Health (ORWH) through the Bench to Bedside Program award #884515.

